# The *UV RESISTANCE LOCUS 8*-Mediated UV-B Response Is Required Alongside *CRYPTOCHROME1* For Plant Survival Under Sunlight In The Field

**DOI:** 10.1101/2021.12.08.471623

**Authors:** Reinhold Stockenhuber, Reiko Akiyama, Nicolas Tissot, Misako Yamazaki, Michele Wyler, Adriana B. Arongaus, Roman Podolec, Yasuhiro Sato, Stefan Milosavljevic, Alex Widmer, Roman Ulm, Kentaro K. Shimizu

**Author notes:** These authors contributed equally to this work (Reiko Akiyama, Nicolas Tissot). Corresponding author (KKS), E-Mail, (RU).

## Abstract

As sessile organisms, plants are subjected to fluctuating sunlight including potentially detrimental ultraviolet-B radiation (UV-B). In *Arabidopsis thaliana*, experiments under controlled conditions have shown that *UV RESISTANCE LOCUS 8* (*UVR8*) controls photomorphogenic responses for acclimation and tolerance to UV-B; however, its long-term impacts on plant performance remain poorly understood in naturally fluctuating environments. Here we quantified the survival and reproduction of different *Arabidopsis* mutant genotypes in diverse field and laboratory conditions. We found that *uvr8* mutants produced more fruits than wild type in growth chambers with artificial low UV-B conditions but not in natural field conditions. Importantly, independent double mutants of *UVR8* and the blue-light photoreceptor gene *CRYPTOCHROME 1* (*CRY1*) in two genetic backgrounds showed a drastic reduction in fitness in the field. UV-B attenuation experiments in field conditions and supplemental UV-B in growth chambers demonstrated that UV-B caused the conditional *cry1 uvr8* lethality phenotype. RNA sequencing in different conditions revealed a large number of genes with statistical interaction of *UVR8* and *CRY1* mutations in the presence of UV-B in the field. Among them, Gene Ontology analysis identified enrichment of categories related to UV-B response, oxidative stress, photoprotection and DNA damage repair. Our study demonstrates the functional importance of the *UVR8*-mediated response across life stages *in natura*, which is partially redundant with *CRY1*, and provides an integral picture of gene expression associated with plant environmental responses under diverse environmental conditions.

## Introduction

Plants have to be able to cope with changing environments to survive and reproduce. Field studies uncovered that the function of few or even single genes can affect fitness components, namely biomass and fruit production (Kerwin et al., 2017; Külheim et al., 2002; Taylor et al., 2019; Tian et al., 2003). Recently, a growing number of studies have shown that plant gene expression patterns and phenotypes observed in the laboratory are often different from those in natural environments (Kerwin et al., 2017; Kudoh, 2016; Sato et al., 2019b; Shimizu et al., 2011; Song et al., 2018; Yamasaki et al., 2017). One environmental factor involved in such difference is light condition. Photoreceptor-mediated perception and responses to solar radiation contribute to plant survival and reproduction in the field (Galen et al., 2004; Liu et al., 2004; Mazza and Ballaré, 2015; Moriconi et al., 2018; Rai et al., 2019; Sellaro et al., 2019; Yankovsky et al., 1995), thereby providing a key to understand plant adaptation to naturally fluctuating environments.

*Arabidopsis thaliana* (*Arabidopsis*) has distinct gene families encoding photoreceptors that sense the light environment. A total of thirteen photoreceptors from five distinct gene families are known in *Arabidopsis*, namely five red/far-red light-perceiving phytochromes (phyA-E); seven blue/UV-A photoreceptors, comprising two cryptochromes (cry1 and cry2), three Zeitlupe family members (ztl, fkf1, and lkp2), and two phototropins (phot1 and phot2); and the UV-B photoreceptor UV RESISTANCE LOCUS 8 (UVR8) (Galvão and Fankhauser, 2015; Podolec et al., 2021a; Rizzini et al., 2011). UV-B is a potentially damaging abiotic stress factor that may affect survival, and thus the fitness and distribution of plant populations (Demarsy et al., 2018; Escobar-Bravo et al., 2017; Jenkins, 2017). Importantly, UVR8 orchestrates UV-B-induced photomorphogenesis and stress acclimation in plants. The fundamental role of UVR8 was shown in both controlled chamber conditions and sun simulators that mimic natural sunlight, in which pronounced adverse effects on the phenotypes of *uvr8* mutants and their survival were observed (Brown et al., 2005; Favory et al., 2009; Kliebenstein et al., 2002). In contrast, *Arabidopsis* plants defective in *UVR8* grown in the field did not show higher mortality at seedling stage or an obvious aberrant morphology though they did display reduced photoprotective pigment levels (Coffey et al., 2017; Morales et al., 2013). Despite recent progress in understanding its molecular mechanism (Podolec et al., 2021a), effects of the *UVR8* gene on plant fitness are still rather unclear under field conditions.

Recent studies demonstrate overlapping signalling mechanisms and partially redundant functions of UVR8 with other photoreceptors, especially with cry1 (Lau et al., 2019; Podolec and Ulm, 2018; Ponnu et al., 2019; Rai et al., 2020, 2019; Tissot and Ulm, 2020; Wang and Lin, 2020). Some of these studies showed a short-term influence of *UVR8, CRY1* and *CRY2* genes on plant growth and gene expression profiles under sunlight, encompassing seedling growth within a month (Rai et al., 2019) or transcriptional changes after a short exposure to sunlight (Rai et al., 2020). However, little is known about long-term impacts of these genes on plant fitness and gene expression profiles. For an *in natura* understanding of photoreceptors it is necessary to quantify plant fitness and gene expression under various environmental conditions.

In this study, we investigated plant survival and reproduction of *uvr8* mutants as well as the effects of a potential overlap of photoreceptor function of UVR8 with cry1 under diverse field and laboratory conditions. Furthermore, we conducted RNA-seq to examine gene expression changes among the field and laboratory conditions. By quantitatively assessing fitness components and underlying molecular mechanisms of different mutants in various environments, we addressed the following questions:

1. Are fitness components (i.e., survival and reproductive output) associated with the *UVR8-*mediated response?
2. Do *cry1* and *uvr8* mutations have synergistic effects on fitness in the field?
3. Which genes show interaction effects between *cry1* and *uvr8* mutations in their expression in the field?

## Results

### Fitness reduction is associated with UVR8-mediated UV-B response detected in growth-chamber conditions

To examine how the UVR8-mediated UV-B response affected fitness components, three independent *uvr8* null mutants and their respective wild types *(uvr8-1* in the L*er* background, *uvr8-7* in Ws, and *uvr8-19* in Col-0; Table S1) were grown in growth chambers and the reproductive output and growth of plants of each genotype were analyzed (Tables S2 – S4). The growth chambers had constant, low levels of UV-B supplied with fluorescent white-light tubes (Chamber-UV), providing approximately 1.5% of the daily UV-B present in the field in summer (Table S5). The *uvr8* mutants produced significantly more fruits than the wild types (Chi-squared = 80.66, p < 2.20*10^-16^, Fig. 1; Chi-squared = 23.37, p < 1.43*10^-6^, Fig. S1A). Because no significant differences in fruit length (Chi-squared = 0.35, p = 0.532, Fig. S1B) or seeds per fruit (Chi-squared = 0.31, p = 0.578, Fig. S1C) were observed, this indicates that the per-plant seed number was increased. Similarly, the *uvr8* mutants produced more overall biomass (fresh weight), an indicator of growth, than the wild types (Chi-squared = 6.00, p = 0.014, Fig. S1D). By contrast, no significant differences in either reproductive output or plant growth were detected under UV-B-exclusion conditions (Chi-squared = 1.64, p = 0.201, Fig. S1E). Taken together, these results suggest that a significant reduction in fitness is associated with the response to UV-B mediated by functional alleles of the *UVR8* gene in the Chamber-UV condition.

**Figure 1.**
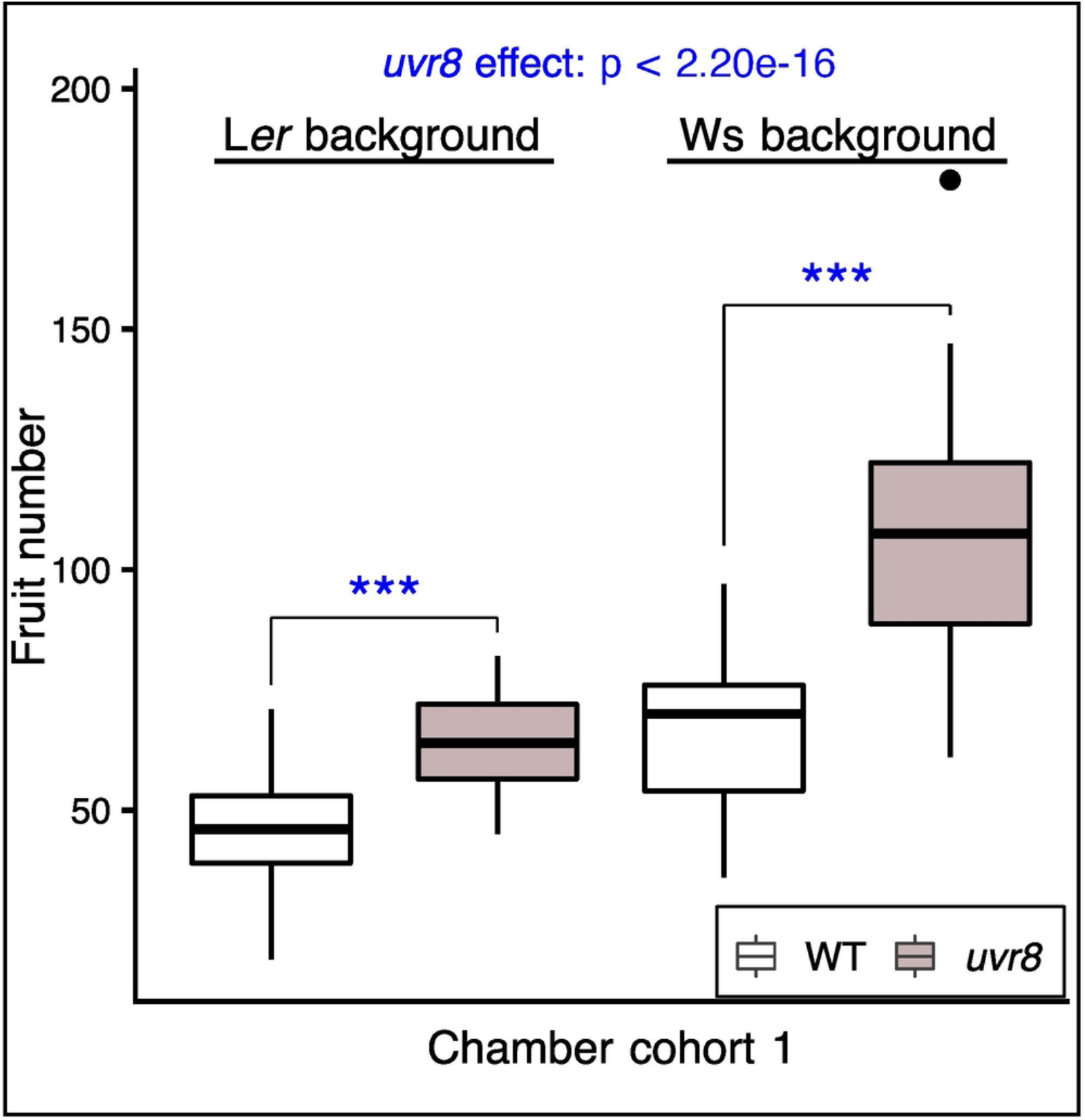

### Performance of single *uvr8* mutants in diverse field conditions

We conducted experiments in field conditions in Zurich, a location representing the natural range of *Arabidopsis*, to investigate whether *uvr8* mutants show reduced fitness in sunlight. To schedule experiments, we determined the *Arabidopsis* growth season from herbarium specimen and field observations. Among 116 specimens collected in or near the Kanton Zurich deposited in the Zurich herbaria, nine had flowers and/or fruits in July and August (Fig. S2A), whereas others flowered in early spring, indicating overwintering. Field observations in Zurich also showed that *Arabidopsis* bore flowers and fruits throughout spring and summer in addition to the overwintering cohorts (Sato et al., 2019a). Thus, we studied both overwintering and non-overwintering cohorts. The UV-B dose during growth of the non-overwintering cohorts was several times higher than that of the overwintering cohorts (Table S5). No significant differences in fitness components (fruit number or survival; Fig. S2B and S2C) or plant habit (Fig. S2d) were observed between *uvr8-1* mutant and L*er* wild type in various field experiments in Zurich (Table S3), regardless of season or developmental stage.

We also grew plants at a high-elevation site in the Swiss Alps (Mountain cohort). In this environmental condition, *uvr8*-*1* showed a higher mortality in comparison to the L*er* wild type in a non-overwintering cohort (Chi-squared = 9.29, p = 0.002, Fig. S2B). Unfortunately, other fitness traits could not be assessed due to massive herbivory damage that occurred after bolting on all plants. The maximum UV-B irradiation was higher in the Mountain cohort than in Zurich (Table S5).

### Double mutants of *UVR8* and *CRY1* show severe defects in field conditions

To test for potential functional redundancy between UVR8 and cry1, we grew wild type, *cry1, uvr8*, and *cry1 uvr8* plants (in two different backgrounds, L*er* and Col; Table S1) in two seasons. Our statistical analysis was centered on whether *cry1 uvr8* exhibited more severe defects in fitness components than the addition of the single mutant defects would explain. This was tested by including an interaction term (statistical interaction) between *uvr8* (functional or non-functional) and *cry1* (functional or non-functional), integrating data of the two backgrounds (Tables S3 and S4). We examined an overwintering cohort, in which plants were exposed to field conditions from the seed stage (Fig. 2, Overwintering cohort 3). We found that *cry1 uvr8* plants strongly reduced the seedling establishment (statistical interaction, *cry1* x *uvr8*, Chi-squared = 10.75, p = 0.001, incorporating both accessions, the same below; Fig. 2A) and growth (Fig. 2B). A large part of the leaves of the double mutants showed yellowing and eventually chlorosis (white arrows in Fig. 2B). After 123 days (4 months), surviving plants of this genotype had severely reduced biomass (*cry1* x *uvr8*, Chi-squared = 16.25, p = 5.54*10^-5^, Fig. S2E), whereas after 132 days, all *cry1 uvr8* plants were dead (Fig. S2F) without developing fruits *(cry1* x *uvr8*, Chi-squared = 303.10, p < 2.20*10^-16^, Fig. 2C). These fitness component data suggest that negative effects associated with the double mutants are synergistic rather than additive. We also conducted an experiment in a non-overwintering condition, where only a small number of replicates was measurable due to a dysfunction in the irrigation systems. Nonetheless, the interaction effect on inflorescence dry weight was similarly significant despite small sample numbers *(cry1* x *uvr8*, Chi-squared = 15.49, p = 8.28 *10^-5^, Fig. S3A right panel). After 45 days in field conditions, *cry1 uvr8* double mutants of Col background were visibly different (Fig. S2G), and a significant interaction of *cry1* and *uvr8* effects on anthocyanin content was detected *(cry1* x *uvr8*, Chi-squared = 30.99, p = 2.59*10^-8^, Fig. S2G).

**Figure 2.**
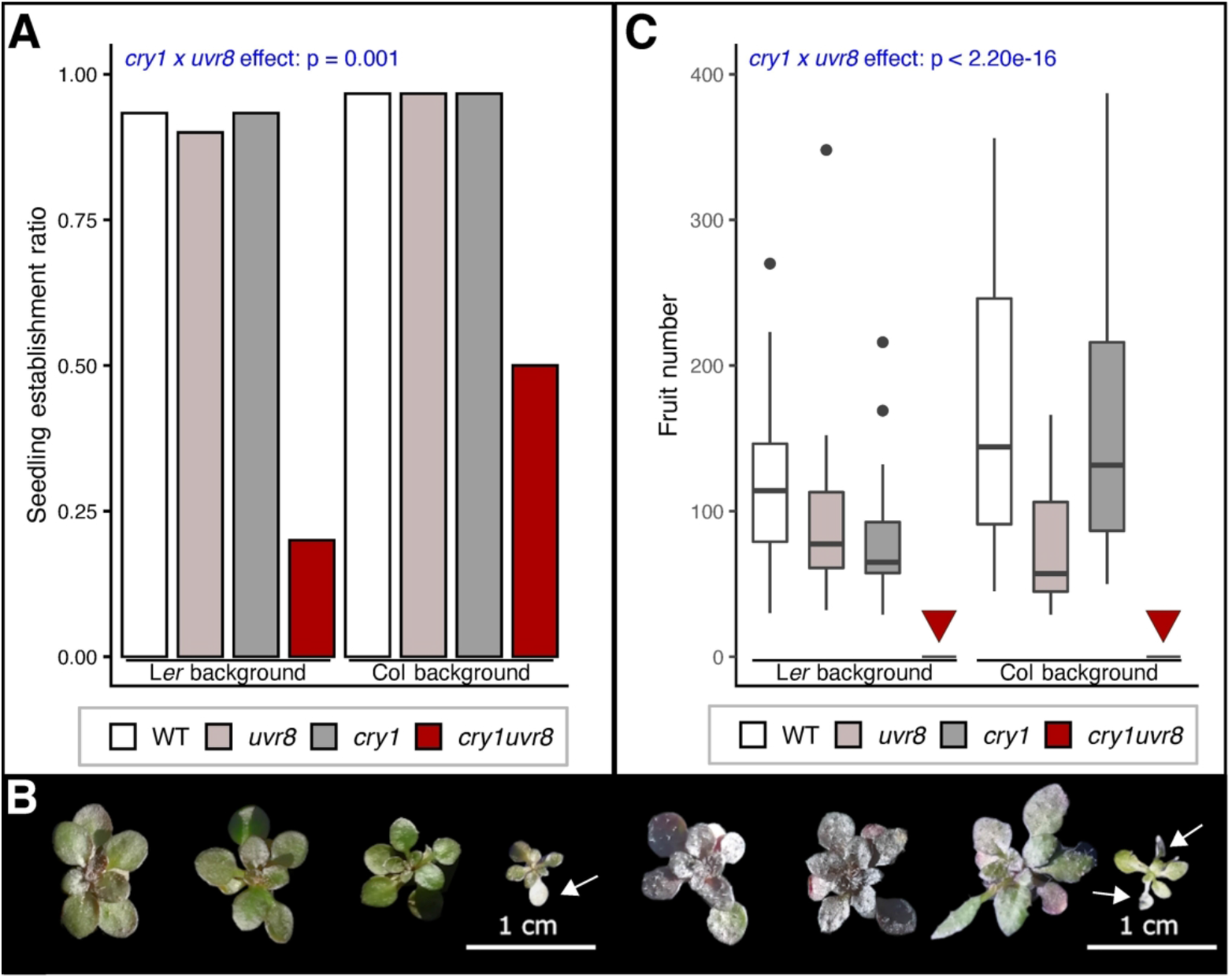

### UV causes the conditional *cry1 uvr8* lethality

We next studied whether UV irradiation is responsible for the severe defects of *cry1 uvr8* double mutants by growing plants in experimentally manipulated UV levels in field. We constructed two types of UV-screening tents with different levels of UV filtering (Fig. 3A). The first type was covered by a UV-absorbing filter film (Rosco #226) and termed Low-UV, which transmitted approximately 1% of UV-B irradiation (Table S5). The second type was covered by a more transparent UV filter film (Rosco #130) and termed UV-med, which transmitted approximately 25% of the daily UV-B dose in July, similar to the daily UV-B dose in winter conditions (Table S5). Fig. 3B illustrates the natural fluctuation of UV-B intensity, temperature, and relative humidity under UV-med and Low-UV (e.g., days 2 and 3 were rainy with lower UV-B intensity). These data show that temperature and relative humidity are comparable, but the degree of UV-B level is different between the two experimental conditions (Fig. 3B, Table S5).

**Figure 3.**
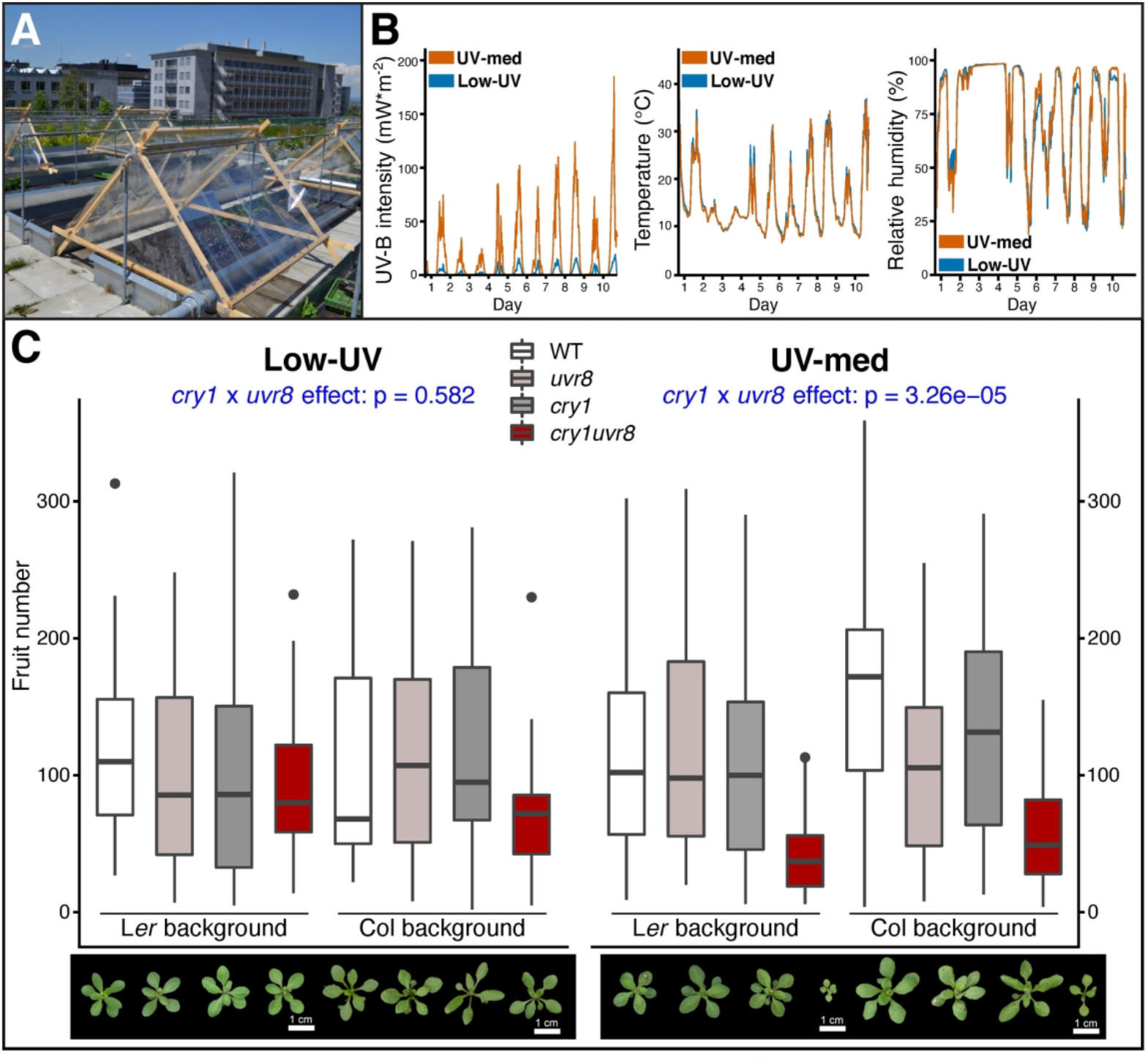

We grew a non-overwintering cohort from seeds for a complete life cycle in the field (Fig. 3C and D; Table S2; Non-overwintering cohort 2). Two quantitative fitness components related to seed production were measured, *i*.*e*., fruit number and inflorescence dry weight, which were highly correlated (adjusted R^2^ ≥ 0.869; Fig. S3B). In UV-med, *cry1 uvr8* double mutants showed reduced growth (Fig. 3C, right panel); however, they did not die prematurely, which enabled tissue sampling for subsequent RNA extraction and expression analysis (see below). Importantly, in Low-UV, *cry1 uvr8* growth was comparable to wild type (Fig 3C, left panel), thus demonstrating that solar UV is responsible for the *cry1 uvr8* defects. Similar to the previous field experiments, we statistically tested the interaction effect of *UVR8* and *CRY1* on fitness components (Fig. 3C, Fig. S3A and C, Table S3 and S4), which was significant in UV-med (fruit number: Chi-squared = 17.26, p = 3.26*10^-5^; inflorescence dry weight: Chi-squared = 13.25, p = 2.73*10^-4^) but not in Low-UV (fruit number: Chi-squared = 0.30, p = 0.582; inflorescence dry weight: Chi-squared = 0.07, p = 0.790). These results suggest that the synergistic defect of *UVR8* and *CRY1* is only detectable in the presence of considerable UV irradiation. An effect of UV-B on fitness was additionally supported by a significant three-way interaction effect of *UVR8, CRY1*, and UV-B condition (ANOVA; three-way interaction; fruit number Chi-squared = 4.35, p = 0.037; inflorescence dry weight Chi-squared = 4.33, p = 0.038). In addition, a similar pattern was observed in another small-scale non-overwintering cohort (Non-overwintering cohort 3, Fig. S3a).

These UV attenuation experiments corroborated that UV is the causal factor for the growth defects in *cry1 uvr8* plants. Indeed, *cry1 uvr8* but not the respective single mutants showed high mortality (statistical interaction Chi-squared = 114.09, p < 2.2*10^-16^) and strongly impaired growth in laboratory conditions with supplemental UV-B specifically (Table S5; Fig. S4a-c), in agreement with previous findings in L*er* background (Tissot and Ulm, 2020). Combined with results from the UV attenuation experiments in the field, these data suggest that the reduced fitness of *cry1 uvr8* double mutants in the field is attributable to UV-B exposure.

### Gene expression profiles among field and laboratory conditions corresponds to fitness component results

We performed a transcriptome analysis to characterize potential statistical interaction effects of cry1 and UVR8 on gene expression profiles, similar to the fitness components described above. We obtained RNA-seq data of 36 seedling samples in total representing three biological replicates each of four L*er* genotypes (L*er*, *hy4-2.23N, uvr8-1*, and *hy4-2.23N uvr8-1*; note that *hy4-2.23N* is a *cry1* mutant allele) grown in three conditions (UV-med and Low-UV in the Non-overwintering cohort 2, and UV-B-exclusion in the chamber cohort 6). We performed a principal component analysis (PCA) to examine the most influential factors on gene expression. The first two axes (PC1 and PC2) had a major effect on gene expression (46.1% and 13.6%, respectively, Fig. 4a). PC1 corresponds to the difference between the field and chamber conditions, supporting a major difference between regulated and field conditions. PC2 corresponds to the genotypic differences, driven by the separation of double mutants in UV-med. Consistent with PC2, the number of genes with statistically significant interaction of *uvr8* and *cry1* mutations (fdr-adjusted p ≤ 0.05) was highest in UV-med (1,438 genes; Table S6), much lower in Low-UV (141 genes; Table S7), and very low in the UV-B-exclusion condition (5 genes; Table S8), supporting a synergy of UVR8 and cry1 in response to UV irradiation.

**Figure 4.**
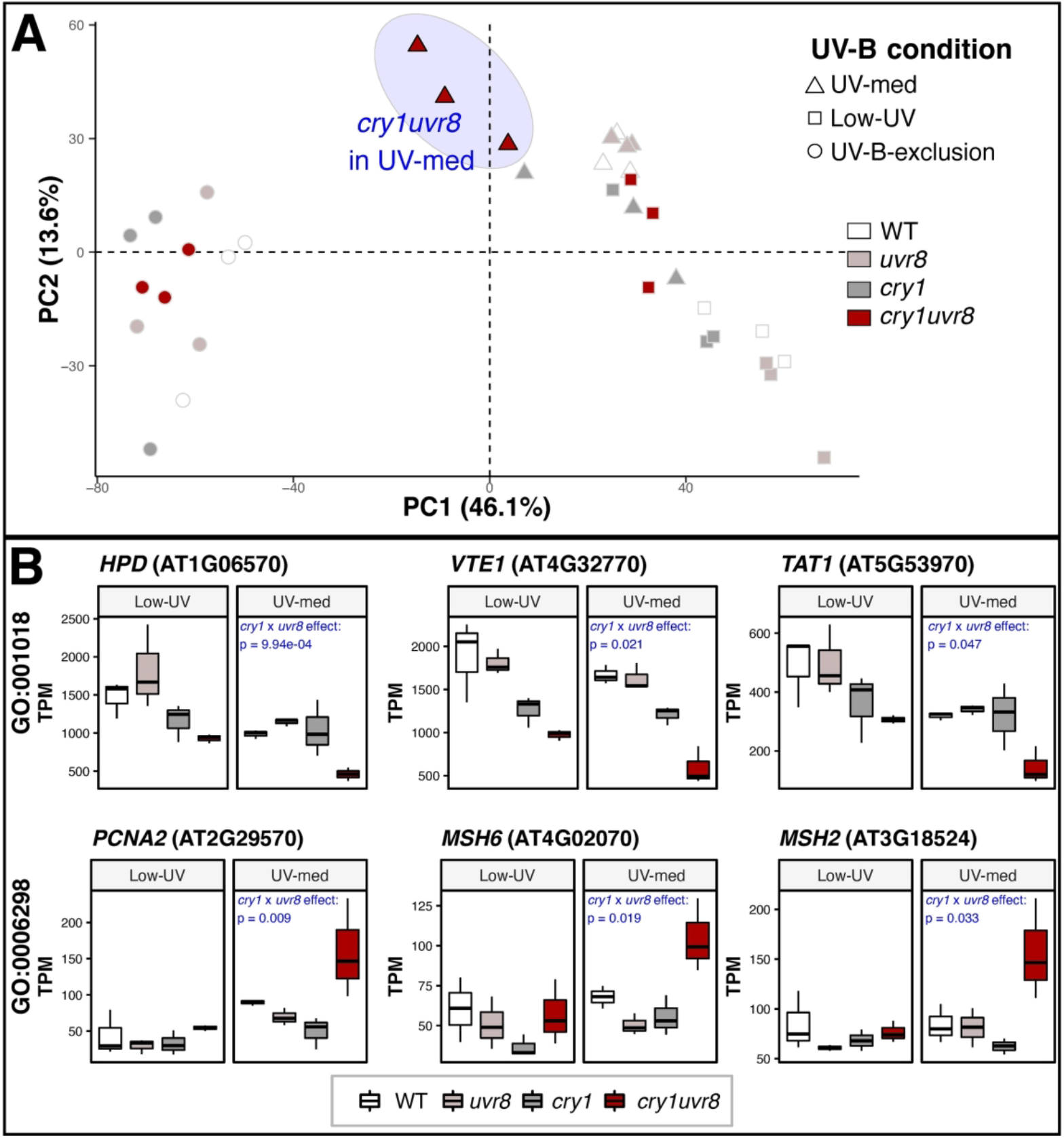

We further examined genes with significant interaction in UV-med. Of these, 513 and 520 genes showed >2-fold decrease or increase in expression, respectively. Among the 513 genes with reduced expression, 239 Gene Ontology (GO) terms were significantly enriched (Table S9). Response to light stimulus (GO:0009416) and specifically response to UV-B (GO:0010224) and blue light signaling (GO:0009785) were enriched, along with pathways that are directly affected by cry1 and UVR8 function, e.g., flavonoid biosynthetic process (GO:0009813) and regulation of photomorphogenesis (GO:0010099). Categories related to photosynthesis and response to oxidative stress were enriched, including response to oxidative stress (GO:0006979) and vitamin E biosynthetic process (GO:0010189). Next, among the 520 genes with increased expression, 180 GO categories were enriched (Table S10). Interestingly, DNA damage repair terms were enriched, e.g., mismatch repair (GO:0006298), double strand break repair (GO:0006302), or recombinational repair (GO:0000725). Fig. 4B illustrates the decreased and increased expression of the double mutants in UV-med (right panels) and the non-significant difference in expression of the same genes in Low-UV (left panels), respectively.

## Discussion

### Fitness consequence of the *UVR8*-mediated responses

*Arabidopsis* studies of stress responses have shown that mutant plants lacking stress resistance sometimes grow faster or reproduce more than wild types in the absence of the stressor. Such an intrinsic cost is known in single-gene mutants regarding disease (Tian et al., 2003), herbivory (Sato et al., 2019a; Züst et al., 2011), and herbicide resistance (Purrington and Bergelson, 1999; Roux et al., 2004) in *A. thaliana*. For example, *csr1-1* and *ixr1-2* mutants, which gain resistance to acetolactate synthase or cellulose synthase inhibitors, exhibit 36-44% reduction in total silique production in the absence of herbicide (Roux et al., 2004), suggesting negative pleiotropic effects due to the alteration of physiological process. Similar to these findings, we report here the intrinsic cost of a response pathway resulting in abiotic stress tolerance, the UVR8-mediated UV-B response. The absolute value of its fitness cost is comparable to previous studies and depended on measured fitness components and on growth conditions owing to the inducible nature of the response. In the UV-B-exclusion conditions, no significant differences in fitness components of wild types and *uvr8* mutants of different *Arabidopsis* backgrounds were detected, suggesting negligible induction of UV-B responses. Differences were pronounced in conditions with UV-B levels corresponding to approximately 1.5% of natural UV-B irradiation, but in the presence of a more significant stressor, i.e. sunlight in Zurich, no significant difference was detected. Furthermore, in contrast to naturally occurring mutations in herbicide resistance and herbivore defense, *UVR8* is generally highly conserved (Fernández et al., 2016). We did observe lethality of *uvr8* mutants in a small-scale experiment at an alpine site. We would like to note that the data alone cannot establish if UV-B irradiation was directly responsible for the lethality observed in the Mountain cohort. The total amount of UV irradiation in the alpine environment is subject to large fluctuations and may not have been greater than that in Zurich (Blumthaler, 2012). In addition altitudinal gradients in irradiation, temperature, rainfall, and others are correlated (Körner, 2003). Therefore, larger scale studies encompassing diverse environments would be recommended to assess a broader picture of the *UVR8* gene function *in natura*.

### Synergistic effect of *CRY1* and *UVR8* mutations on fitness components shown by statistical interactions

The interaction in the downstream cascade of different photoreceptors is of major interest in photobiology. Recent studies suggested a common mechanism of UVR8- and cryptochrome-mediated inhibition of COP1 (Favory et al., 2009; Lau et al., 2019; Podolec et al., 2021a; Podolec and Ulm, 2018; Ponnu et al., 2019), and a *cry1 cry2 uvr8* triple mutant showed lethality under UV-B in natural conditions (Rai et al., 2019). By growing single and double mutants in diverse experimental settings, we here provide substantial evidence that functional alleles of *UVR8* and *CRY1* synergistically prevent severe defects under UV-B in field conditions.

We found parallel evidence for fitness increase/decrease among different background accessions with single or double mutation on *UVR8* and *CRY1*. The consistent results under diverse environmental conditions strongly suggest roles of these two genes in plant adaptation to sunlight *in natura*. Significant statistical interaction indicated that the loss-of-function effects of *UVR8* and *CRY1* on fitness components were synergistic. In addition, when double mutants were grown from seeds, complete synthetic lethality was observed. Similar results were obtained in both overwintering and non-overwintering cohorts in Zurich. The use of UV filters in the field restored normal *cry1 uvr8* plant growth, further confirming that their defects are caused by UV, in agreement with a previous study of *cry1 cry2 uvr8* triple mutants in the field (Rai et al., 2019). Moreover, the growth of plants in a chamber with supplemental UV-B recapitulated the elevated sensitivity of *cry1 uvr8* double mutants alleles used in this work when compared to the respective single mutants and wild type, in agreement with previous findings (Tissot and Ulm, 2020).

Cryptochromes evolved before the split of plants and animals and may have played an ancestral role for short-wavelength sensing and response. *UVR8* originated in green algae and its function was shown to be conserved up to land plants (Allorent et al., 2016; Fernández et al., 2016; Podolec et al., 2021a; Rizzini et al., 2011; Tilbrook et al., 2016). Our data supports that *UVR8* and *CRY1* are synergistically required for plant survival under sunlight (Rai et al., 2019; Tissot and Ulm, 2020).

### Synergistic effect of *CRY1* and *UVR8* mutations on the expression of genes responsible for UV response and DNA integrity

Similar to fitness related traits, we detected significant statistical interaction of *uvr8* and *cry1* mutations at the the gene expression level. Among the genes with reduced expression, GO analysis suggested biological processes of mainly three groups: light response, photosynthesis and oxidative stress. Light response related GOs were consistent with previous field studies that performed pairwise comparisons of mutant to wild type genotypes (Morales et al., 2013; Rai et al., 2020). Among these terms were response to UVB (GO:0010224), blue light signaling (GO:0009785), flavonoid biosynthesis (GO:0009813), and regulation of photomorphogenesis (GO:0010099). The photosynthetic machinery has also been previously shown to be susceptible to high light stress, and especially UV (Demarsy et al., 2018; Lütz and Seidlitz, 2012; Takahashi et al., 2010). An important role in maintaining its function under such conditions is attributed to certain protective compounds, such as tocopherol (Vitamin E), and their defect can lead to photooxidation and chlorosis (Havaux et al., 2005; Ksas et al., 2015; Miret and Munné-Bosch, 2015). Consistent with chlorotic leaves found in double mutants (Fig. 2b), the expression levels of genes involved in Vitamin E biosynthesis (Fig. 4B upper panels, Table S11) showed significant statistical interaction and were also reduced in *cry1 uvr8* double mutants.

The genes that showed increased gene expression for the statistical interaction in UV-med were mainly enriched for methylation and DNA repair related terms, such as mismatch repair (GO:0006298). The upregulation of two DNA mismatch repair protein genes *(MSH2* and *MSH6*) and the *PCNA2* gene (Fig. 4B lower panels, Table S11) attributes a potential role to UVR8 and cry1 in the maintenance of DNA integrity under solar UV(-B). We speculate that the impaired UV responses described above resulted in DNA damage by UV irradiation, and thus DNA repair pathways may have been upregulated to compensate the damages.

### Conclusion

Our data highlights the complex nature of light responses throughout plant life stages and the importance of combining field and laboratory experiments. By using a genetically tractable species such as *Arabidopsis* we were able to add to the understanding of the molecular bases of abiotic stress responses in plants. To test the ecological relevance of cry1 and UVR8, we applied a dual method: assessment of the quantitative fitness and gene expression variation of different genotypes, including single and double mutants in a factorial design. This approach enabled us to gain insight on the interaction effects of two important photoreceptors perceiving blue light and UV-B, respectively, crucial for UV-B tolerance in the field. Thus, our study showcases the value of combining mutant analyses with ecological functional genomics in understanding the molecular basis of plant environmental response *in natura* (Rai et al., 2021).

## Materials and Methods

### Genotype information

*Arabidopsis thaliana* mutants *uvr8-1* (Kliebenstein et al., 2002), *hy4-2.23N* (Ahmad and Cashmore, 1993), and *hy4-2.23N uvr8-1* (Tissot and Ulm, 2020) are in the Landsberg *erecta* (L*er*) background; *uvr8-19* (Podolec et al., 2021b), *cry1-304* (Mockler et al., 1999), and *cry1-304 uvr8-19* (Podolec et al., 2021b) in Columbia-0 (Col-0); and *uvr8-7* (Favory et al., 2009) in Wassilewskija (Ws). See Table S1 for further details on these lines.

### Experimental conditions

We prepared common gardens in Zurich at the Irchel Campus of the University of Zurich (47°23’46.1” N, 8°33’04” E, 500 m altitude) and Calanda, Grisons (46°53’16.1” N, 9°29’21.6” E, 2000 m altitude, Mountain cohort), as well as growth chambers. The Calanda site was kindly provided by the Gemeinde Haldenstein (Switzerland) and managed and permitted by the Plant Ecological Genetics Group in the Institute of Integrative Biology of the ETH Zurich (Switzerland).

For chamber experiments, we used custom-built growth chambers (Kälte 3000) equipped with Regent “Easy 5 Cool White” (FDH-39W) and Regent “!GroLux” (FDH-39W) batten luminaires in a 2:1 ratio. We used two different conditions for experiments in growth chambers. Long-day conditions with 16-h light (22°C, 60% rH, 120–160 μE light) and 8-h darkness (20°C, 60% rH) was used for plant pre-treatment as well as in Chamber-UV, UV-B-exclusion, and +UV-B conditions. Short-day conditions were 8-h light (18°C, 60% rH, 120–160 μE light) and 16-h darkness (16°C, 60% rH).

Each growth experiment was with multiple genotypes arranged in random block design. Table S2 shows the experimental duration, the plant growth stage at transfer for each experiment, the initial number of transferred individuals per genotype, the UV-B levels, whether the experiment was disrupted before data could be acquired and the corresponding figures results are displayed in. Setup for growth chamber experiments closely resembled that for the field experiments, with the addition of mild insecticide (50 g/l RAVANE 50, Schneiter AGRO) and fungicide treatments (1 g/l Thiovit Jet, Maag Garden) every 14 days.

Environmental data were recorded using UV-Microlog miniature data loggers (sglux). These loggers are weather-resistant and provide logging function of three independent environmental variables over longer time intervals. The loggers were equipped with a UV-B diode (TOCON_E_OEM, sglux), and an integrated temperature and external humidity sensor (rH in percent). The output shown in this study for the UV-Microlog is the erythemally weighed UV-B intensity in mW*m^-2^.

The loggers were used to record environmental data for several days, performed at least once for all of the field and chamber conditions. In Non-overwintering cohort 2, three loggers were used in parallel to compare UV attenuation with unfiltered UV-B levels (Table S5). The data loggers were placed on even ground directly in the compartments.

### Plant material

Plants were transferred to the field either as seeds or as seedlings at the five-leaf rosette stage, respectively (Table S2). We directly transferred seeds to the field in both cohorts to investigate full life cycles, although experiments were occasionally destroyed owing to the vulnerability of early seedling stages by natural fluctuations such as heavy rain or drought effects. Seedlings therefore were brought to the field for some of the experients, as is commonly done in *Arabidopsis* field studies (Sato et al., 2019a; Tian et al., 2003) as a backup for measurements in case seed-derived plants were lost.

For the seed stage, we stratified seeds by putting them on 0.8% agar plates with 1/2 Murashige & Skoog (MS) medium or in Eppendorf tubes with tap water at 4°C in darkness for up to 72 h. Three to five seeds were then transferred to the surface of watered standard soil (Einheitserde) in plastic pots (8 x 8 x 7 cm). The pots were kept at room temperature overnight and then transferred to the field. After germination and before the leaves of plants growing in the same pot began to touch, thinning was performed in all experiments by cutting off and removing above-ground plant parts in order to obtain one plant per pot.

As preparations for the transfer of plants to the field at five-leaf stage, seeds were sown on 0.8% agar plates with 1/2 MS medium. After 72 h at 4°C in darkness, plates were transferred to growth chambers with long-day conditions to induce germination. Germinated plants were then transferred to soil (Einheitserde) in plastic pots (8 x 8 x 7 cm) and kept growth chambers with short-day condition to avoid early flowering onset until plants reached a five-leaf stage. For acclimation, the seedlings in the plastic pots were transferred to shaded places in the common garden 24–48 h before placing them in the compartments.

Plants for growth chamber experiments were prepared in the same way as plants for field experiments transferred at seed stage until potting. The potted plants were then placed in one of the chamber conditions (Chamber-UV, UV-B-exclusion, and +UV-B). placed in growth chambers for experiments in chamber conditions (Chamber-UV, UV-B-exclusion, and +UV-B).

### Field site growth experiments

In the Zurich common garden, each compartment was filled with a 15-cm layer of Rasenerde (Top-Dressing) and enclosed by a slug barrier. We watered each compartment automatically (three fine-spraying valves per compartment, 10 minutes duration, set at 05:00 and 21:00, respectively) between March and November and manually between November and March when the water supply was turned off to avoid frost damage to the water supply system. Pots were arranged at least 10 cm from the edges of the compartments and distributed with at least a 2-cm gap between pots.

To test the influence of UV on mutant lines of *UVR8* and *CRY1*, we prepared a total of six tents with wooden frames covered with UV-blocking (Rosco #226) or -transmitting (Rosco #130) filter (n=3 for each filter type). Both filter types are recommended (Aphalo et al., 2012) for photobiological experiments and are commonly used (e.g., Morales et al., 2013; Rai et al., 2020). The experiments were conducted in non-overwintering cohorts to avoid breakage and snow cover of UV filters by winter conditions.

At the high-elevation field site (Mountain cohort, 2000m), a 1 x 2 m compartment without enclosure was prepared. Ten centimeters of the top soil layer was exchanged with standard soil (Top-Dressing Rasenerde) and metal wire and fleece were embedded 10 cm below the soil surface of the compartment to avoid disturbances by fossorial animals.

### Chamber growth experiments

In the chamber experiments, we used the normal Chamber-UV (under fluorescent lamps, as described above). In addition, UV-B-exclusion conditions were established by applying UV-blocking filter film (Rosco #226, S4 Fig d) and supplemental UV-B in +UV-B was added from Philips “TL20W/01RS” narrowband UV-B tubes. Pots were placed in the corresponding chamber conditions and watered manually every 2–3 days. Water levels were controlled to be at ca. 1.5-cm height after watering. During flowering, plant floral stems were bound to a wooden stick in the center of the pot to avoid contact between flower stems from different individuals.

### Plant trait measurements

Throughout the experiments, a number of different plant traits were assessed. We measured fitness components survival and reproductive output (fruit number, inflorescence dry weight; and in Chamber cohort 2, fruit length and seeds per fruit were additionally assessed). Furthermore, biomass (fresh weight of aboveground plant parts) in Overwintering cohort 3 as well as in Chamber cohort 4 was assessed. In Non-overwintering cohort 3 anthocyanin accumulation was measured. Survival was recorded at the time of harvest as the presence/absence of a plant in each pot. In addition, seedling establishment (after germination success) was measured in Overwintering cohort 3 to infer the survival of germinated plants. These individuals were then monitored for their survival until flowering onset.

To assess biomass, rosettes were harvested and immediately placed in liquid nitrogen to avoid drying. After collection, the frozen plant material was weighed on a precision balance.

Reproductive output in chamber experiments was evaluated by counting the fruit number (siliques). Fruit number was assessed in plants of Chamber cohorts 1–3, Overwintering cohort 1, and Overwintering cohort 3 as well as Non-overwintering cohort 1-2. In addition, in Non-overwintering cohort 2-3, aerial plant parts above rosette leaves were harvested together after primary and secondary inflorescences ceased flower production, dried at 60°C for at least 24 h, and then weighed on a precision balance to determine inflorescence dry weight. Inflorescence dry weight and the fruit number were highly correlated (Fig. S3B) and therefore only the former was measured for Non-overwintering cohort 3, in which the plants grew large and the fruit number was very high. Absence of plants at time of harvest was recorded as zero count and statistical analysis was performed without zeros for a more conservative analysis, except for fruit number in the case of Overwintering cohort 3, where we performed analysis with zeros, as no surviving double-mutant plants were observed (see statistical analysis).

### Transcriptomic analysis

#### RNA extraction

Field samples of the L*er* genotypes were collected following 2 weeks (14 d) growth in field (UV-med and Low-UV in Non-overwintering cohort 2) and chamber (UV-B-Exclusion in Chamber cohort 6) conditions. We sampled rosette leaves between 11:00 and 14:00 to avoid gene expression fluctuation caused by the effects of diurnal rhythms. Material was collected in 1.5-ml vials and directly transferred to liquid nitrogen, and then stored at −80°C until RNA extraction. We used the RNeasy Plant Mini Kit (Qiagen) for RNA extraction following the manufacturer’s protocol without DNase treatment. RNA quantity was measured using a Qubit 2.0 (Thermofisher Scientific) and then diluted to 25 ng/μl per sample.

#### Library preparation and sequencing

Total RNA (500 ng per sample) was used to synthesize libraries using a TruSeq Stranded mRNA Kit v2 (Illumina). Cluster generation was performed at the Functional Genomics Center Zurich (FGCZ, Switzerland) using 10 mM of pooled normalized libraries on the cBOT with a TruSeq PE Cluster Kit v3-cBot-HS (Illumina). Subsequently Illumina HiSeq 4000 sequencing was performed to generate the reads.

#### Data processing

The data analysis framework SUSHI (Hatakeyama et al., 2016) was employed to process raw reads. Standard settings implemented in SUSHI were used for RNA-seq data processing. Data analysis was performed according to with the following steps:

For quality analysis, *FastQC* v 0.11 (Andrews, 2010) was used. To align the reads to the Araport 11 *Arabidopsis* reference genome (Cheng et al., 2017), *STAR* (Dobin et al., 2013) was used. We then assigned mapped reads to genomic features with FeatureCounts and used CountQC, implemented in Qualimap (García-Alcalde et al., 2012) for quality control after counting.

Further analysis was performed in RStudio v1.0.143 implementing R v 3.3.3 and above (http://www.r-project.org/). Packages ggplot2 (Wickham, 2016) and ggpubr (Kassambara, 2019) were used to create graphics. Mapped and quantified reads were used for a principal component analysis (PCA) on all genes to identify the most contributing dimensions, employing packages DEseq2 (Love et al., 2014), factoextra (Kassambara and Mundt, 2020) and FactoMineR (Lê et al., 2008).

Differential gene expression analysis was conducted with DESeq2. Our goal was to identify gene-gene interaction effects within and across UV-attenuation treatments.

To estimate statistical power, we fitted two fully factorial models. Model 1 included factors for gene function of *CRY1* and *UVR8*, treatment and statistically significant interactions within and between genes and treatment. We increased statistical power by reducing complexity in Model 2, which was based on by-treatment subsets of data, separating Low-UV from UV-med. GO enrichment analysis was performed on this set of genes with topGO (Alexa and Rahnenführer, 2021) using the elim algorithm. To reduce redundancy due to GO term hierarchy, the identified GO categories were filtered to those categories with at least ten and less than 1000 annotations.

### Relative anthocyanin accumulation

Leaf anthocyanin content was determined by spectrophotometry according to an adjusted method from Schmidt & Mohr (Schmidt and Mohr, 1981). Pre-weighed *Arabidopsis* leaf tissue was placed in 800 μL extraction buffer (2-propanol:HCl:H2O in 18:1:81 percent by volume), boiled for 3 min and then kept at room temperature in darkness overnight. The samples were then centrifuged at 10’000 rpm for 2 min and the absorption of extracted anthocyanin was measured at 535 nm and 650 nm. Leaf anthocyanin content was then calculated by subtracting the absorption at 650 nm from that at 535 nm and dividing this by fresh weight [y = (A_535_ - A_650_)/fresh weight].

### Statistical analysis

All statistical analyses were performed in R v 3.3.3 and above (http://www.r-project.org/). In box plots, bars indicate the median, boxes indicate the interquartile range. Whiskers extend to the most extreme data point that is no more than 1.5 times the interquartile range from the box, and outliers are indicated with dots.

Explanatory variables consisted of *uvr8*_mutant (fixed effect), *cry1*_mutant (fixed effect), treatment (fixed effect), background (random effect), and block (random effect). The variables *uvr8*_mutant and *cry1*_mutant indicate gene functions. For each variable, whether the following mutations *uvr8-1, uvr8-7, uvr8-19, hy4-2.23N, cry1-304, hy4-2.23N uvr8-1*, or *cry1-304 uvr8-19* existed or not was scored as yes or no. See Table S12 for an overview of scoring. Treatment was defined only for UV manipulation experiments in the field. This variable consisted of two levels created by the different filter types: UV-med or Low-UV. Background consisted either of L*er*, Col, or Ws. Within each block, one individual of a combination of *uvr8*_mutant, *cry1*_mutant, treatment, and background was assigned, except for Overwintering cohort 1, Overwintering cohort 2, Non-overwintering cohort 1, and Mountain cohort, where a random design across each compartment was applied.

Using these variables, we built different models depending on the purpose and set-up of the experiment. To test the effect of mutant genotype on fitness components in chamber experiments, we included *uvr8*_mutant as explanatory variables in the model. In chamber and field experiments using *cry1*_mutant, we built a model for each of the summer and winter cohorts with *uvr8*_mutant and *cry1*_mutant as explanatory variables in order to test the effect of mutants on fitness components. In field experiments, we also examined the effect of the interaction between *uvr8*_mutant and *cry1*_mutant by adding an interaction term in the model. To test the effects of mutants, UV, and interaction thereof on fitness components in the UV manipulation experiment in field, we built a model with *uvr8*_mutant, *cry1*_mutant, treatment (referring to the UV-conditions Low-UV and UV-med), two-way interactions (*uvr8*_mutant x *cry1*_mutant, *uvr8*_mutant x treatment, and *cry1*_mutant x treatment), and a three-way interaction (*uvr8*_mutant x *cry1*_mutant x treatment) as explanatory variables in the model. In all models, we included background and block as explanatory variables, when applicable.

We adopted a linear model framework that was suitable for binary, count, and continuous traits with additional sources of trait variation considered (Faraway, 2016). Survival data were scored binary, and therefore generalized linear models (GLM) or generalized linear mixed models (GLMM) with binomial distribution were built. In the case of Overwintering cohort 3, data showed complete separation, *i.e*. the range of values of a response variable for one group of an explanatory variable did not overlap with that of an(other) group(s) of the same explanatory variable. In this case, no model would fit the data properly. Therefore, we transformed the data by adding a count of one (1) to each individual observation before fitting a model.

For non-survival data (i.e., biomass, anthocyanin, fruit number, inflorescence dry weight, the length of fruit, and the number of seeds per fruit), we used linear models (LM) and linear-mixed models (LMM) when normality could be assumed by histograms and univariate Shapiro-Wilk normality tests. Otherwise, we used GLM or GLMM. We used R packages *stats, glmmTMB v 0.2.3* (Bolker et al., n.d.), and *lme4 v 1.1* (Bates et al., 2014).

Analysis with GLM and GLMM was done in the following steps. First, we built three models with different distributions, i.e., Poisson, negative binomial, as well as quasi-Poisson (or the type-I). Models that failed to converge or that converged with warnings were excluded from further steps. When at least two models were applicable, we determined the best model using the AICtab function of the package *bblme v1.0.20*, the model with the lowest likelihood ratio score was considered the best model. The best model was then examined for the fit of data by visually examining simulated standard residual plots (≥ 250 simulations per model to reduce stochastic errors), by one-sample Kolmogorov-Smirnov test and by outlier test using the package *dHARMA v0.2.3* (Hartig, 2018). When the best model did not appropriately fit the results, we built new models with Poisson, negative binomial, or quasi-Poisson distribution with zero-inflation parameters using the package *glmmTMB*, with zero-inflation parameters applying to all observations (zi=~1 or zi=~.) or absences varying by specific factors (e.g. treatment, gene functions, see Supplemental R-scripts). These models were evaluated using the same steps as above. In the case of LM and LMM, we generally built a single model including all random factors and then continued with examining the best model for the fit of data as above.

Once the best model was identified, we tested the significance of the fixed effect(s) on the response variable by conducting an analysis-of-variance using the function Anova in the package *car v3.0-2*. When one or more interaction terms was present in the model, we used a type-III Wald Chi-squared test. Otherwise, we used a type-II Wald Chi-squared test (Table S3).

## Data availability

Data availability: Raw sequence reads used for RNA-seq analysis in this study are available at the DNA databank of Japan (SAMD00199169 - 646SAMD00199240). Source data files to the statistical analysis are available in the provided data package ().

## Acknowledgments

We thank Ueli Grossniklaus, Angela Hancock, Masaomi Hatakeyama, Takato Imaizumi, Akira Nagatani, Christoph Ringli, and Takashi Tsuchimatsu for helpful discussion, and Carlos D. Crocco for early contributions to the project. We further thank the Plant Ecology Group (PEG) of the ETH Zurich, especially Jörg Leuenberger, Regina Zaech, and Marc-Jacques Maechler, for organizing and maintaining the high elevation site at Grisons; Reto Nyffeler and staff at the Zurich Herbaria (Z+ZT) for their kind support in the herbaria study; the URPP Global Change and Biodiversity; Giulia Ghielmetti and Andreas Hueni from the Department of Geography for supplying field equipment and environment data; and Lucas Mohn and Aki Morishima from the Department of Evolutionary Biology and Environmental Studies for their help throughout the experiments.

